# Protein mismatches caused by reassortment influence functions of the reovirus capsid

**DOI:** 10.1101/322651

**Authors:** Deepti Thete, Pranav Danthi

## Abstract

Following attachment to host receptors via σ1, reovirus particles are endocytosed and disassembled to generate infectious subvirion particles (ISVPs). ISVPs undergo conformational changes to form ISVP*, releasing σ1 and membrane-targeting peptides from the viral μ1 protein. ISVP* formation is required for delivery of the viral core into the cytoplasm for replication. We characterized the properties of T3D^F^/T3D^C^S1, a S1 gene monoreassortant between two laboratory isolates of prototype reovirus strain T3D: T3D^F^ and T3D^C^. T3D^F^/T3D^C^S1 is poorly infectious. This deficiency is a consequence of inefficient encapsidation of S1-encoded σ1 on T3D^F^/T3D^C^S1 virions. Additionally, in comparison to T3D^F^, T3D^F^/T3D^C^S1 undergoes ISVP-to-ISVP* conversion more readily, revealing an unexpected role for σ1 in regulating ISVP* formation. The σ1 protein is held within turrets formed by the λ2 protein. To test if the altered properties of T3D^F^/T3D^C^S1 are due to a mismatch between σ1 and λ2 proteins from T3D^F^ and T3D^C^, properties of T3D^F^/T3D^C^L2 and T3D^F^/T3D^C^S1L2, which express a T3D^C^-derived λ2, were compared. The presence of T3D^C^ λ2 allowed more efficient σ1 incorporation, producing particles that exhibit T3D^F^-like infectivity. In comparison to T3D^F^, T3D^F^/T3D^C^L2 prematurely converts to ISVP* uncovering a role for λ2 in regulating ISVP* formation. Importantly, a virus with matching σ1 and λ2 displayed a more regulated conversion to ISVP* than either T3D^F^/T3D^C^S1 or T3D^F^/T3D^C^L2. In addition to identifying new regulators of ISVP* formation, our results highlight that protein mismatches produced by reassortment can alter virus assembly and thereby influence subsequent functions of the virus capsid.

## IMPORTANCE

Cells coinfected with viruses that possess a multipartite or segmented genome reassort to produce progeny viruses that contain a combination of gene segments from each parent. Reassortment places new pairs of genes together generating viruses in which mismatched proteins must function together. To test if such forced pairing of proteins that form the virus shell or capsid alters the function of the capsid, we investigated properties of reovirus variants in which the σ1 attachment protein and the λ2 protein that anchors σ1 on the particle, are mismatched. Our studies demonstrate that a σ1-λ2 mismatch produces particles with lower level of encapsidated σ1, consequently decreasing virus attachment and infectivity. The mismatch between σ1 and λ2 also altered the capacity of the viral capsid to undergo conformational changes required for cell entry. These studies reveal new functions of reovirus capsid proteins, and illuminate both predictable and novel implications of reassortment.

## INTRODUCTION

For segmented RNA viruses such as influenza virus, rotavirus, and bluetongue virus, genetic reassortment is a crucial evolutionary mechanism (1). Reassortment of gene segments allows viruses to acquire novel genetic markers that are necessary to overcome host defenses (2). Reassortment can also lead to emergence of strains with superior replicative fitness or those with a capacity to infect new hosts (3). One well studied segmented RNA virus that undergoes reassortment, both in cell culture and in infected animals, is mammalian orthoreovirus (reovirus) (4, 5). The double stranded RNA genome of reovirus consists of 10 segments. Reassortants formed by coinfection of prototype reovirus strains T1L and T3D has led to the identification of determinants that control efficiency of viral replication in cell culture and those that influence viral pathogenesis in a mouse model (6–12). As with other segmented viruses (13–18), reovirus reassortment is not random and certain gene combinations are favored whereas others are excluded (5, 19). Though the mechanisms driving preferred selection of parental alleles are still not fully elucidated, one factor that governs the generation and recovery of a reassortant is the maintenance of functional protein-protein interactions in the capsid proteins or replication machinery of the virus (1).

The genome segments of reovirus are encapsidated within two concentric protein shells: the core and the outer capsid (20). During replication of the virus, viral mRNAs are packaged within the core along with viral polymerase and NTPase (λ3 and μ2 respectively)(20). The core is a turreted icosahedron formed from λ1, λ2 and σ2 (21). Pentamers of the λ2 protein form the turret and this structure can only be formed in the presence of λ1 and σ2 ((22, 23). μ1 and σ3, major components of the outer capsid form heterohexamers (24) . The core is connected to the μ1-σ3 layer via interaction of μ1 with both σ2 and λ2 (25, (26). Trimers of σ1 are anchored within the turret by the flap domains of λ2 (21, 25). The minor outer-capsid protein σ1 is only found in particles that also contain σ3 and μ1 in its native state (27–29). This effect may be a consequence of changes in the conformation of the λ2 flap due to the presence of the adjacent μ1 protein (26, 30, 31). Fully formed virions initiate infection of new host cells and this process requires the function of multiple capsid proteins. Attachment of host cells occurs via the σ1 protein (32). Proteolytic degradation of σ3 within the endosome results in formation of ISVPs (33). ISVPs undergo further conformational changes to form ISVP*s, resulting in changes to the λ2 and μ1 structures that facilitate the release of σ1 trimers and μ1-derived peptides (27, 29). The μ1 peptides facilitate the delivery of cores into the cytoplasm (29, 34, 35). Conformational changes in the capsid also activate the viral transcriptional machinery, which allows expression of the viral mRNA and completion of the remainder of the viral replication cycle (27). Analogous to other viral systems, it is anticipated that the protein-protein interactions that govern the proper assembly of stable virus capsids influence properties of the virus capsid.

Two laboratory isolates of prototype reovirus strain T3D, T3D^F^ (F, Fields) and T3D^C^ (C, Cashdollar) display differences in their infectious properties, including in the morphology of viral replication factories, cell killing capacity, and in vivo replication efficiency (36–38). Here, we characterized the properties of capsids of T3D^F^ and T3D^F^/T3D^C^S1, a monoreassortant bearing the S1 gene from T3D^C^ in an otherwise T3D^F^ virus. We found that in comparison to T3D^F^, particles of T3D^F^/T3D^C^S1 display an assembly defect, encapsidating less σ1. Particles of T3D^F^/T3D^C^S1 therefore exhibit a diminished capacity to attach and infect cells. Surprisingly, in comparison to T3D^F^, capsids of T3D^F^/T3D^C^S1 undergo conformational changes characteristic of ISVP-to-ISVP* conversion without an appropriate trigger. The effects of T3D^C^S1 on the attachment and ISVP* conversion efficiency of T3D^F^ could be overcome by introduction of a matched λ2-encoding T3D^C^ L2 gene. In addition to highlighting changes in σ1 that influence its encapsidation, these studies identify a previously unknown role for σ1 and λ2 in controlling conformational changes required for cell entry. These findings provide new insights into understanding how interaction and matches between proteins that form viral capsids influence properties of the capsid and may influence the generation or replicative capacity of reassortant viruses.

## RESULTS

### The infectivity of T3D^F^ is compromised by introduction of the T3D^C^ σ1 protein

A single gene reassortant between prototype reovirus strains T1L and T3D, which contains the μ1-encoding M2 gene segment from T3D in an otherwise T1L genetic background exhibits enhanced attachment to host cells (39). Reovirus attachment is a function of the σ protein (32, 40). The μ1 protein does not make physical contact with σ1 and therefore the effect of μ1 on σ1 function is unexpected (26, 39, 41). Curiously, the μ1 proteins of T1L and T3D display ~ 98% identity with the two proteins differing in only 15 out of 708 residues, which are scattered throughout the primary sequence of the protein (42). Thus, it appears that even a minimal difference in the properties of analogous proteins from two different parents can influence the phenotype of reassortant progeny. To determine whether this unforeseen phenotype of reassortment extends to other gene combinations and other virus strains, we characterized the properties of T3D^F^/T3D^C^S1, an S1 gene monoreassortant between two laboratory isolates of strain T3D: T3D^F^ (F, Fields) and T3D^C^ (C, Cashdollar). The S1 gene reassortant T3D^F^/T3D^C^S1 is ideal since unlike prototype reovirus strains such as T1L, T2J, and T3D, where the S1 gene sequences are highly divergent, the S1 genes of T3D^F^ and T3D^C^ differ minimally (36). The S1 gene encodes two proteins from overlapping reading frames, σ1 and σ1s (43, 44). The σ1 proteins of T3D^F^ and T3D^C^ differ at amino acid residues 22 and 408, resulting in a Valine-to-Alanine change at residue 22 and a Threonine-to-Alanine change at residue 408 (36). Because the 5’ end of S1 that produces a polymorphism in σ1 at position 22 also encodes σ1s in an alternate reading frame, it results in a Glutamine-to-Histidine change at residue 3 in σ1s (36).

During initial characterization of T3D^F^ and T3D^F^/T3D^C^S1, we observed that in comparison to the parental strain T3D^F^, T3D^F^/T3D^C^S1 shows plaques with a predominantly diminished size in L929 cells (Fig 1A). Plaque morphology is a measure of the capacity of the virus to efficiently complete all stages of the viral replication cycle and infect neighboring cells. It is also affected by the host response to virus infection. To compare the capacity of T3D^F^ and T3D^F^/T3D^C^S1 to establish infection in host cells, L929 cells were infected with 3000 virions/cell and relative infectivity was determined at 18 h post infection using quantitative indirect immunofluorescence (45). We observed that in comparison to T3D^F^, T3D^F^/T3D^C^S1 displayed significant lower infectivity (Fig 1B). In some tissues, ISVPs are formed extracellularly (46–48). Under these conditions, infection of host cells is initiated by ISVPs. As observed with the infection initiated by virions, chymotrypsin-generated ISVPs of T3D^F^ infected L929 cells to a greater extent than similarly generated ISVPs of T3D^F^/T3D^C^S1 (Fig 1C). Thus, our data indicate that the presence of T3D^C^S1 gene segment in an otherwise T3D^F^ background produces a virus that has a diminished capacity to establish infection in host cells.

**Figure 1.**
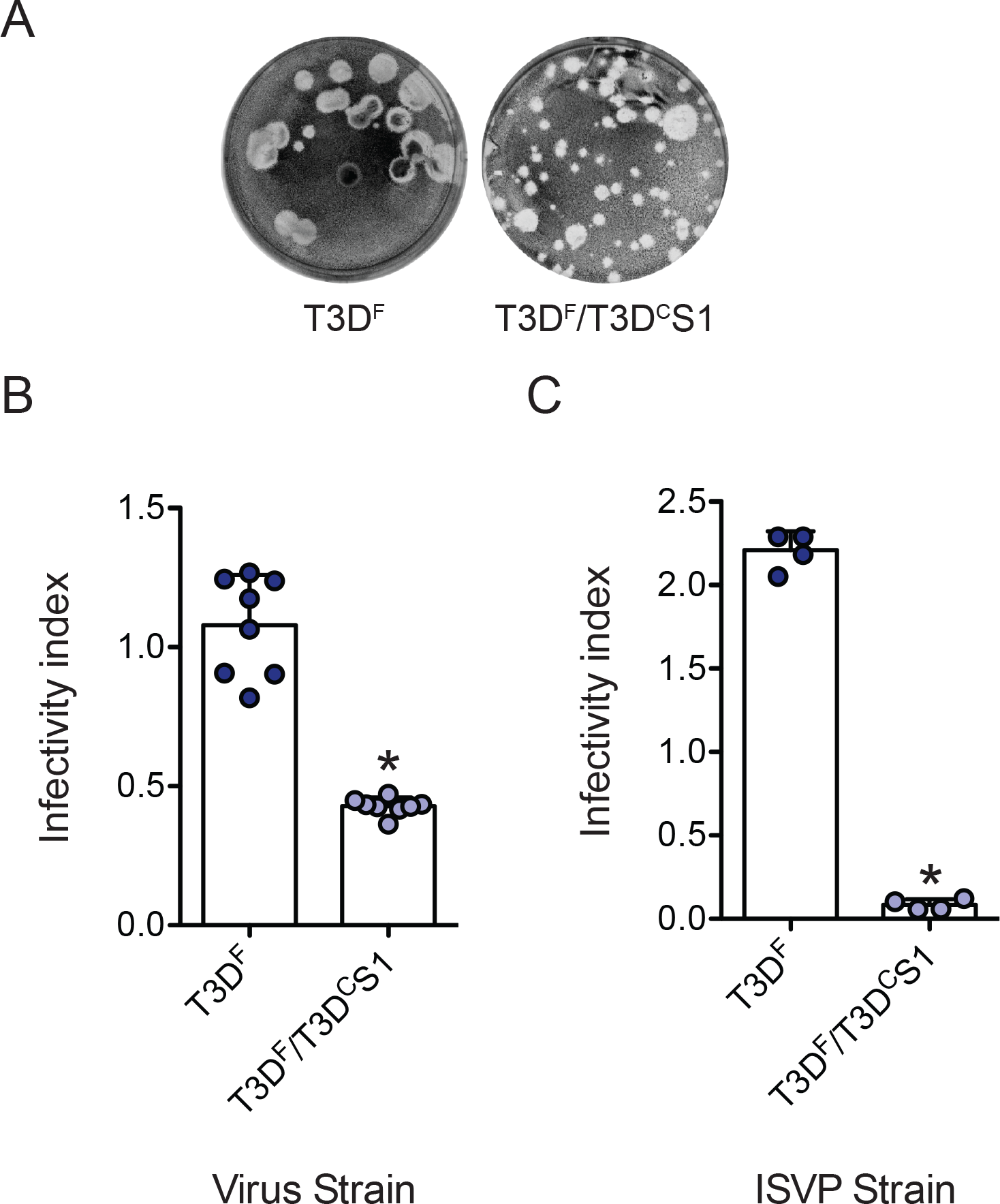
T3D^F^/T3D^C^S1 exhibits small plaque morphology and reduced infectivity in L929 cells. (A) Particles of T3D^F^ or T3D^F^/T3D^C^S1 diluted in PBS were subjected to plaque assay. ATCC L929 cells were adsorbed with (B) 3000 particles/cell of either T3D^F^ or T3D^F^/T3D^C^S1 or (C) 300 ISVPs/cell of either T3D^F^ or T3D^F^/T3D^C^S1 at room temperature for 1 h. (B, C) After incubation at 37°C for 18 h, the cells were subjected to indirect immunofluorescence assay using a LI-COR Odyssey scanner. Relative infectivity was determined by calculating intensity ratios at 800 nm (green fluorescence) representing viral antigen and 700 nm (red fluorescence) representing the cell monolayer. Infectivity index for each independent infection and the sample mean are shown. Error bars indicate SD. *, P < 0.05 as determined by Student’s t-test in comparison to T3D^F^.

### T3D^F^/T3D^C^S1 attaches to host cells less efficiently

The S1 gene segment encoded σ1 protein mediates attachment of type 3 reovirus strains to cell surface receptors (49–51). Since the S1 genes of T3D^F^ and T3D^C^ differ in their primary sequence, we asked if viruses containing each of these proteins differ in their capacity to attach to host cells. We compared the attachment of T3D^F^ and T3D^F^/T3D^C^S1 to adherent L929 cells using a fluorescence-based quantitative binding assay (39). We observed that in comparison to T3D^F^, T3D^F^/T3D^C^S1 bound to cells less efficiently. These data suggest that the presence of T3D^C^ σ1 in the T3D^F^ virus diminishes the capacity of the virus to bind host cells (Fig 2A).

**Figure 2.**
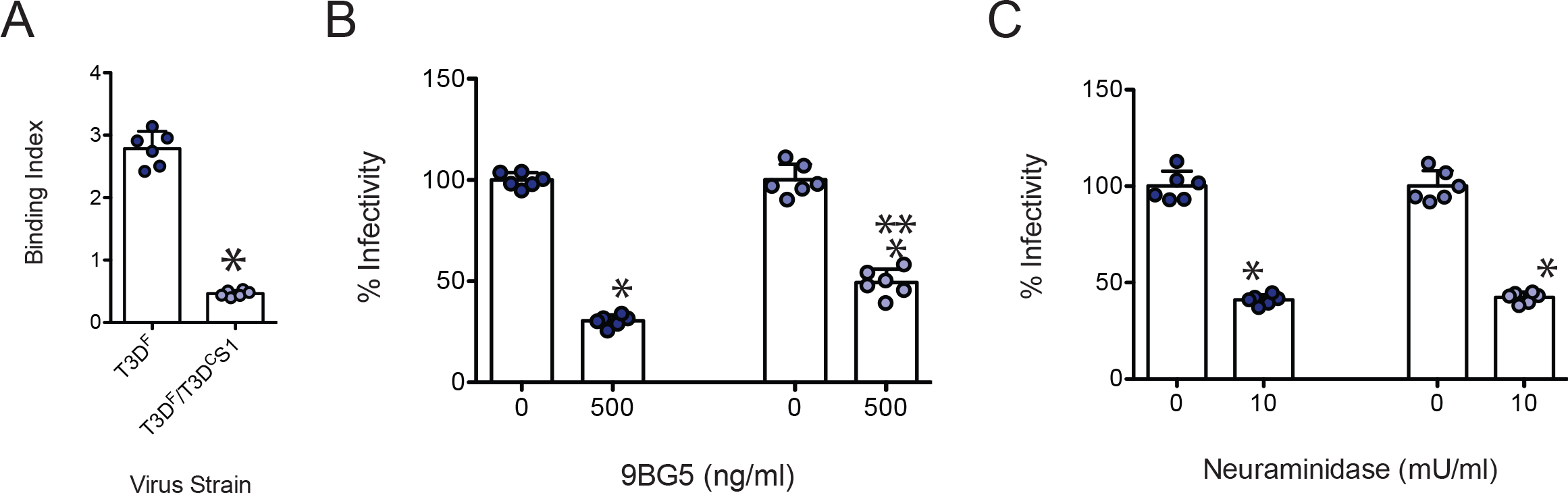
T3D^F^/T3D^C^S1 attaches poorly to L929 cells. (A) Confluent monolayers of L929 cells grown in 96-well plates were adsorbed with 5 × 10^4^ particles/cell of either T3D^F^ or T3D^F^/T3D^C^S1 at 4°C for 1 h. Cell attachment was determined by indirect immunofluorescence of cell-associated particles using a LI-COR Odyssey scanner. Binding index was determined by calculating intensity ratios at 800 nm (green fluorescence) representing viral antigen and 700 nm (red fluorescence) representing the cell monolayer. Binding index for each independent infection and the sample mean are shown. Error bars indicate SD. *, P < 0.05 as determined by Student’s t-test in comparison to T3D^F^ (B) To examine the effect of 9BG5 mAb on virus infectivity, 1 × 10^11^ particles/ml of either T3D^F^ or T3D^F^/T3D^C^S1 were incubated at 4°C with 0 or 500 ng/ml of 9BG5 mAb hybridoma supernatant overnight. This mixture was used to initiate infection of L929 cells in 96-well plates. (C) To examine the role of glycans on infection, L929 cells were pretreated with 0 or 10 mU/ml Neuraminidase. Cells were adsorbed with 3000 particles/cell of either T3D^F^ or T3D^F^/T3D^C^S1 at room temperature for 1 h. (B, C) After incubation at 37°C for 18 h, the cells were subjected to indirect immunofluorescence assay using a LI-COR Odyssey scanner. Relative infectivity was determined by calculating intensity ratios at 800 nm (green fluorescence) representing viral antigen and 700 nm (red fluorescence) representing the cell monolayer. For each virus strain, infectivity index for untreated samples was set to 100%. Percent infectivity for each independent infection and the sample mean are shown. Error bars indicate SD. *, P < 0.05 as determined by Student’s t-test in comparison to T3D^F^. **, P < 0.05 in comparison to similarly treated T3D^F^-infected cells.

The T3D σ1 protein engages JAM-A via its globular head domain, whereas it interacts with sialic acid via regions within the body domain (52, 53). To confirm that both T3D^F^ and T3D^F^/T3D^C^S1 rely on the usual σ1-receptor interactions for type 3 reovirus to attach and infect cells, we assessed the capacity of reagents that diminish interaction of reovirus with each type of receptor to influence the infectivity of T3D^F^ and T3D^F^/T3D^C^S1. Incubation of virions with T3D σ1 head-specific neutralizing monoclonal antibody, 9BG5, which prevents engagement with JAM-A (54, 55), diminished infection by both virus strains (Fig 2B). While we note a small, but statistically significant difference in sensitivity of T3D^F^ and T3D^F^/T3D^C^S1 to 9BG5, we did not further explore this difference for the current study. Pretreatment of cells with *Arthrobacter ureafaciens* neuraminidase to remove cell surface sialic acid also diminished infection by both viruses (Fig 2C). Based on the evidence that both T3D^F^ and T3D^F^/T3D^C^S1 require JAMA and sialic acid to efficiently establish infection in host cells, we think the difference in the attachment and infectivity of T3D^F^ and T3D^F^/T3D^C^S1 are not related to the use of alternate receptors.

### A lower level of σ1 is present on T3D^F^/T3D^C^S1 particles

A reovirus particle can bear a maximum of 12 trimers of the σ1 protein (56). Thus, one possible reason for the lower attachment efficiency of T3D^F^ and T3D^F^/T3D^C^S1 may be lower levels of particle-associated σ1 protein. To test this idea, we compared the level of σ1 relative to another capsid protein, μ1, using quantitative immunoblotting of particles from three independent, freshly CsCl-purified virus preparations. We found that in comparison to T3D^F^, the σ1/μ1 ratio was significantly lower for T3D^F^/T3D^C^S1 (Fig 3A).

**Figure 3.**
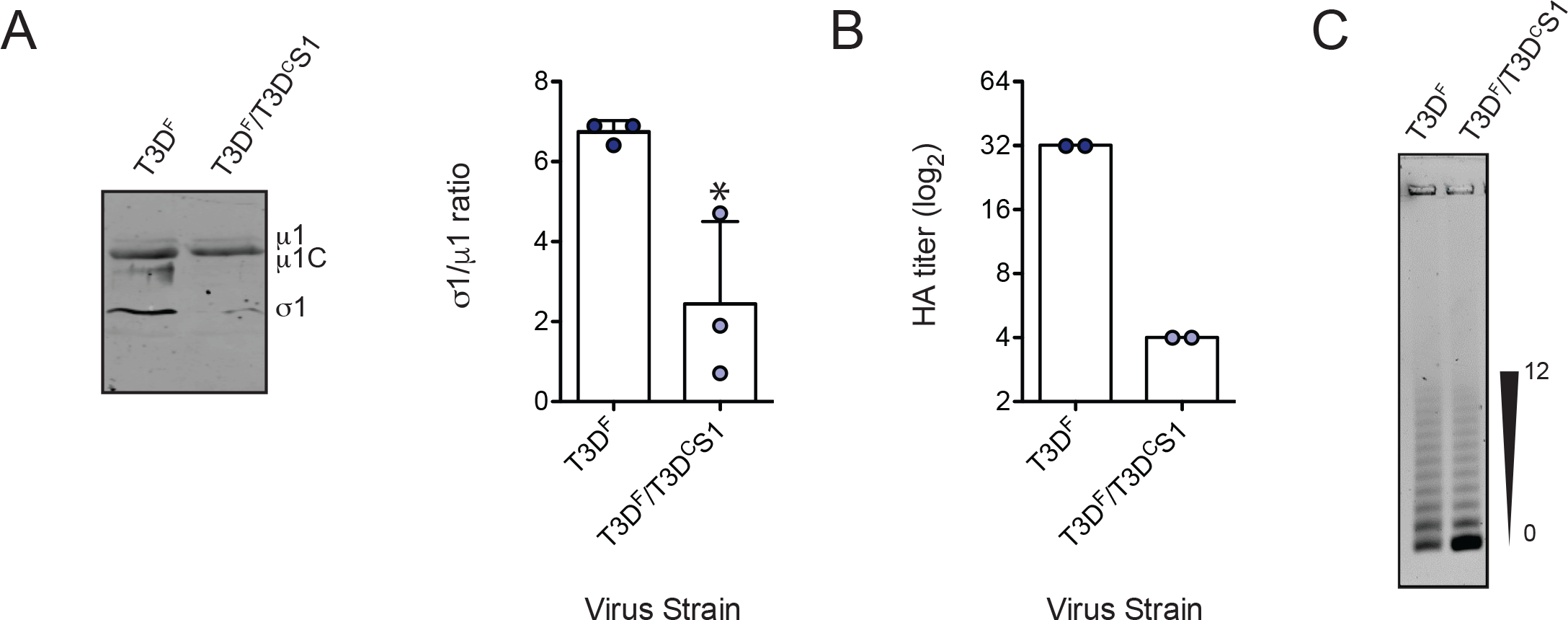
A lower level of σ1 is present on T3D^F^/T3D^C^S1 particles. (A) Purified T3D^F^ or T3D^F^/T3D^C^S1 (2 × 10^10^ particles) virions from 3 independent viral preparations were subjected to immunoblotting using antibodies directed against reovirus p1 protein and T3D σ1 head. Membranes were scanned on a LI-COR Odyssey scanner to determine σ1 and μ1 band intensities. The σ1/μ1 ratios of each independent virus preparation and mean are shown. Error bars indicate SD. *, P < 0.05 as determined by Student’s t-test in comparison to T3D^F^. (B) Purified T3D^F^ or T3D^F^/T3D^C^S1 virions (1 × 10^11^ particles) were serially diluted in PBS and incubated with bovine erythrocytes at 4°C overnight. HA titer expressed as 2 × 10^11^ particles divided by the number of particles per HA unit for each independent sample and the mean value are shown. One HA unit is equal to the number of particles of virus sufficient to produce HA. (C) Virions (1 × 10^11^ particles) were resolved on an agarose gel, stained with colloidal blue staining kit and scanned using a LI-COR Odyssey scanner. The position of particles with lowest and highest number of σ1 trimers is shown.

Type 3 reovirus can agglutinate bovine red blood cells via interaction of σ1 with glycophorins on the surface of cells (57). Hemagglutination capacity can therefore vary with virus concentration. When virus particle concentration is held constant, hemagglutination capacity is indicative of σ1 levels on the particle. We observed that in comparison to T3D^F^, a higher concentration of T3D^F^/T3D^C^S1 was required to agglutinate red blood cells (Fig 3B). These data confirm that in comparison to T3D^F^, particles of T3D^F^/T3D^C^S1 are decorated with lower levels of σ1.

Purified reovirus preparations consist of a conglomeration of virus particles with varying amounts of σ1 trimers (58, 59). The experiments described above (Fig 3A and 3B) cannot reveal if all T3D^F^/T3D^C^S1 particles encapsidate a lower level of σ1 in comparison to T3D^F^ or if a subpopulation of virus particles lack σ1. To distinguish between these possibilities, we took advantage of a method to resolve purified virions on agarose gels based on the level of incorporated σ1 trimers (58). As expected, the preparation of T3D^F^ contained particles bearing a varied number of σ1 molecules. In comparison to T3D^F^, virions of T3D^F^/T3D^C^S1 consisted of higher number of virus particles that encapsidate no σ1 trimers (Fig 3C). The absence of σ1 on a large proportion of T3D^F^/T3D^C^S1 particles likely explains the poor attachment capacity and infectivity of virions of this strain (Fig 1B)

### σ1 is inefficiently encapsidated on T3D^F^/T3D^C^S1 virions

The data presented thus far describe a defect in attachment of T3D^F^/T3D^C^S1 that are produced by changes in properties of the σ1 protein. In addition to the amino acid changes in the σ1 reading frame, polymorphisms in the S1 sequence also result in a change in the σ1s reading frame (36). The nonstructural protein σ1s from T3D^F^ is linked to pathogenesis in vivo but can alter efficiency of viral protein synthesis, host cell cycle, and cell death in cell culture (60, 61). To determine whether differences in the sequence of σ1s in T3D^F^ and T3D^F^/T3D^C^S1 influences viral replication and whether T3D^F^/T3D^C^S1 was compromised at stages of replication other than attachment, we identified conditions under which attachment of T3D^F^ and T3D^F^/T3D^C^S1 were equivalent. We observed that when 2-fold more particles of T3D^F^/T3D^C^S1 were incubated with cells, the attachment of T3D^F^ and T3D^F^/T3D^C^S1 was equivalent (Fig 4A). Correspondingly, when infection was initiated with 2-fold more T3D^F^/T3D^C^S1, the infectivity of T3D^F^ and T3D^F^/T3D^C^S1 was equivalent (Fig 4B). Using 2-fold more T3D^F^/T3D^C^S1, equivalent infectivity of T3D^F^ and T3D^F^/T3D^C^S1 was observed over a wide range of MOIs. These data suggest that when equivalent amount of virus attaches to host cells, other events in the replication cycle that lead to the expression of viral proteins and therefore detection by our quantitative infectivity assay are equivalent.

**Figure 4.**
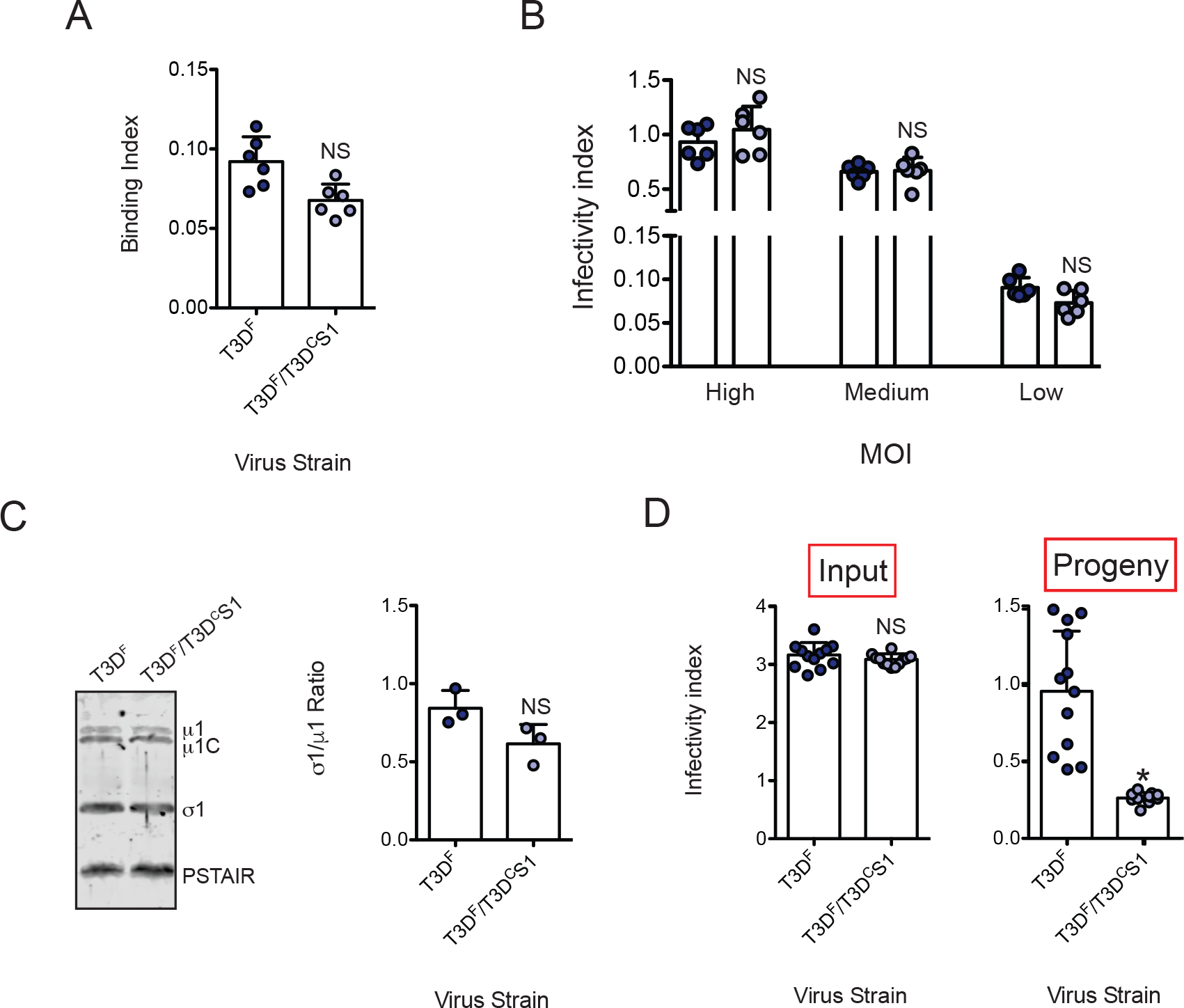
Inefficient assembly contributes to lower level of σ1 on T3D^F^/T3D^C^S1 particles. (A) L929 cells grown in 96-well plates were chilled at 4°C for 15 min and then adsorbed with 5 × 10^4^ particles/cell of T3D^F^ or 1 × 10^5^ particles/cell of T3D^F^/T3D^C^S1 at 4°C for 1 h. Cell attachment was determined by indirect immunofluorescence of cell associated particles using a LI-COR Odyssey scanner. Binding index was determined by calculating intensity ratios at 800 nm (green fluorescence) representing viral antigen and 700 nm (red fluorescence) representing the cell monolayer. Binding index for each independent infection and the sample mean are shown. Error bars indicate SD. *, P < 0.05 as determined by Student’s t-test. (B) ATCC L929 cells were adsorbed with 3000, 300 and 30 particles/cell of either T3D^F^ or 6000, 600 and 60 particles/cell of T3D^F^/T3D^C^S1 at room temperature for 1 h. These conditions are designated as high, medium and low MOI, respectively. After incubation at 37°C for 18 h, the cells were subjected to indirect immunofluorescence assay using a LI-COR Odyssey scanner. Relative infectivity was determined by calculating intensity ratios at 800 nm (green fluorescence) representing viral antigen and 700 nm (red fluorescence) representing the cell monolayer. Infectivity index for each independent infection and the sample mean are shown. Error bars indicate SD. *, P < 0.05 as determined by Student’s t-test. (C) Whole cell lysates of cells infected with 3000 particles/cell T3D^F^ or 6000 particles/cell of T3D^F^/T3D^C^S1 were subjected to immunoblotting using antibodies directed against reovirus protein and T3D σ1 head. Membranes were scanned on a LI-COR Odyssey scanner to determine σ1 and μ1 band intensities. σ1/μ1 ratios for each independent infection and the sample mean are shown. Error bars indicate SD. *, P < 0.05 as determined by Student’s t-test. (D) ATCC L929 cells were adsorbed with 30 particles/cell of T3D^F^ or 60 particles/cell of T3D^F^/T3D^C^S1 at room temperature for 1 h. After incubation at 37°C for 24 h, media supernatant was used to infect fresh L929 cells. Relative infectivity of input virus (in original plate) and that of progeny virus (in fresh plate) was determined by calculating intensity ratios at 800 nm (green fluorescence) representing viral antigen and 700 nm (red fluorescence) representing the cell monolayer. Infectivity index for each independent infection and the sample mean are shown. Error bars indicate SD. *, P < 0.05 as determined by Student’s t-test in comparison to T3D^F^.

Our data indicate that differences in attachment of T3D^F^ and T3D^F^/T3D^C^S1 are due to differences in the levels of particle-associated σ1. Differences in amounts of σ1 on the particle can be either due to changes in steady state levels of σ1 in infected cells or may be due to defective encapsidation or retention of σ1 on virions. To distinguish between these possibilities, we infected cells with equivalent attachment units of T3D^F^ and T3D^F^/T3D^C^S1 (3000 and 6000 particles/cell, respectively) and determined the levels of intracellular σ1 protein by immunoblot analysis. The σ1/μ1 ratio in lysates of cells infected with either T3D^F^ or T3D^F^/T3D^C^S1 was equivalent (Fig 4C). Thus, these data indicate that inefficient encapsidation of σ1 on T3D^F^/T3D^C^S1 particles is not related to poor expression of T3D^C^ σ1 or due to insufficient accumulation of T3D^C^ σ1 in infected cells.

In our experiments shown in Fig 3, we used CsCl-purified viruses to demonstrate that a lower level of σ1 is present on the virus particle. To rule out the possibility that the lower level of particle-associated σ1 on purified particles is a consequence σ1 ejection due to the method of viral purification, we sought to measure the infectivity of unpurified, released viral progeny. Toward this goal, we used an assay recently described to identify host factors required for assembly and release of infectious progeny viruses (62). L929 cells were infected with equivalent attachment units of T3D^F^ or T3D^F^/T3D^C^S1 for 24 h. Consistent with our data shown in Fig 4B, the infectivity index of T3D^F^ and T3D^F^/T3D^C^S1 was equivalent (Fig 4D, input infectivity). The infectivity of the progeny virus produced from this infection was assessed by evaluating the capacity of the released virus present in the media supernatant to establish infection. We found that in comparison to media from T3D^F^ infected cells, the media from cells infected with T3D^F^/T3D^C^S1 showed significantly lower infectivity (Fig 4D, progeny infectivity). These data indicate that assembly and/or release of progeny viruses containing sufficient levels of σ1 is decreased in T3D^F^/T3D^C^S1 even when virions are not subjected to sonication and gradient purification procedures (63). Along with the other data presented above, we think that these findings suggest that in comparison to T3D^F^, particles of T3D^F^/T3D^C^S1 contain a lower level of σ1 due to intracellular defects in σ1 encapsidation.

### T3D^F^/T3D^C^S1 displays increased efficiency of ISVP-to-ISVP* conversion

During cell entry, ISVPs undergo conformational transitions to generate ISVP*s (27). These transitions are likely facilitated by host lipids (64, 65). ISVP-to-ISVP* conversion involves major conformational changes in the particle including (i) reorganization of the μ1 and λ2 proteins; (ii) release of the μ1-derived pore-forming peptides, μ1N and ϕ and (iii) ejection of the σ1 attachment protein (29, (34). If σ1 release is a prerequisite for ISVP-to-ISVP* conversion, a particle with a lower level of encapsidated σ1, or one in which σ1 is held with lower affinity, may be expected to undergo this structural change more readily. To determine if either the amount or the nature of particle-associated σ1 influences the efficiency of ISVP-to-ISVP* conversion, we incubated in vitro generated ISVPs of T3D^F^ and T3D^F^/T3D^C^S1 at increasing temperatures and assessed the trypsin sensitivity of δ fragment as a measure of ISVP* conversion (66). We found that in comparison to T3D^F^, which converts to ISVP*s at 42°C (Fig 5A), T3D^F^/T3D^C^S1 undergoes ISVP-to-ISVP* transition at a significantly lower temperature, ~ 28°C (Fig 5B).

**Figure 5.**
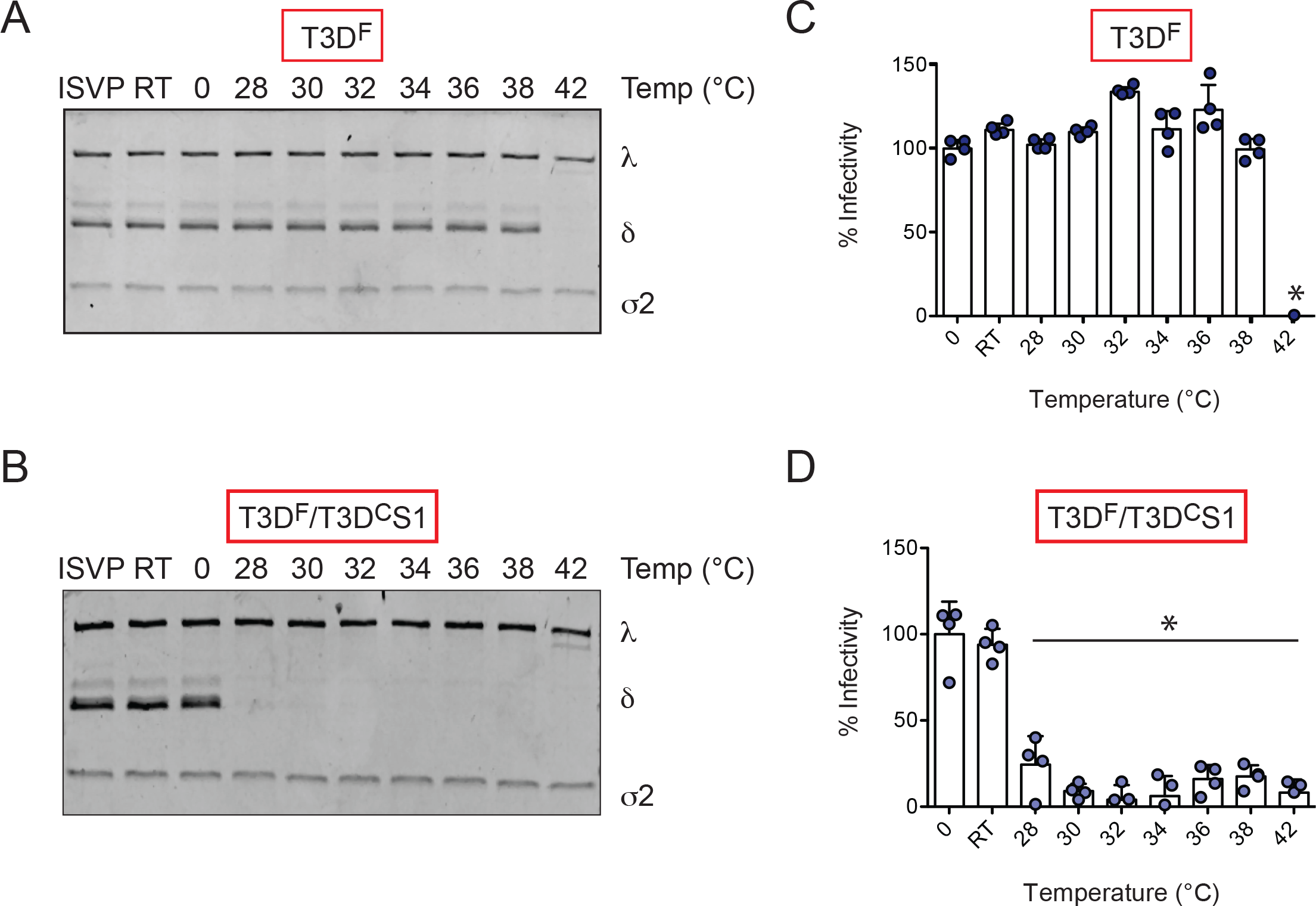
T3D^F^/T3D^C^S1 shows increased efficiency of ISVP-to-ISVP* conversion. (A, B) ISVPs (2 × 10^12^ particles/ml) of T3D^F^ or T3D^F^/T3D^C^S1 were divided into aliquots of equivalent volume and heated at temperatures ranging from 0°C to 42°C for 20 min. The reactions were chilled on ice and digested with 0.10 mg/ml trypsin for 30 min on ice. Following addition of loading dye, the samples were subjected to SDS-PAGE analysis. The gels shown are representative of at least 3 independent experiments. (C, D) ISVPs (2 × 10^12^ particles/ml) of T3D^F^ or T3D^F^/T3D^C^S1 were heated at 0°C to 42°C for 20 min and then used to initiate infection of L929 cells at 300 particles/cell. After incubation at 37°C for 18 h, the cells were subjected to indirect immunofluorescence assay using a LI-COR Odyssey scanner. Relative infectivity was determined by calculating intensity ratios at 800 nm (green fluorescence) representing viral antigen and 700 nm (red fluorescence) representing the cell monolayer. Infectivity index of ISVPs maintained at 0°C was set to 100%. Percent infectivity for each independent infection and the sample mean are shown. Samples with a calculated infectivity below 0% are not shown. Error bars indicate SD. *, P < 0.05 as determined by 1-way ANOVA with Bonferroni’s multiple comparison test in comparison to samples maintained at 0°C.

As a consequence of loss of molecules required for cell entry, ISVP* formation results in diminished infectivity of the particle (67). The temperature at which infectivity is lost therefore serves as a measure of ISVP-to-ISVP* conversion efficiency (65). As an alternative way to assess ISVP* formation, in vitro generated ISVPs of T3D^F^ and T3D^F^/T3D^C^S1 were heated at increasing temperatures and the infectivity of the samples on L929 cells was monitored using the quantitative indirect immunofluorescence assay. We observed that while T3D^F^ lost infectivity at 42°C (Fig 5C), T3D^F^/T3D^C^S1 lost infectivity at a significantly lower temperature (~ 28°C) (Fig 5D). These data indicate that changes to the sequence of the σ1 protein impact the capacity of the particle to undergo ISVP-to-ISVP* transition. These data identify a previously unknown link between the properties of particle-associated σ1 and the propensity for undergoing entry-associated conformational changes.

### Introduction of a matched L2 gene from T3D^C^ restores σ1 encapsidation and infectivity of T3D^F^/T3D^C^S1

The σ1 protein is anchored to the particle within turrets formed by the λ2 protein (26, 68). Thus, the encapsidation or retention of σ1 on the particle can be affected by the nature of the σ1-λ2 interaction. The outer surface of the λ2 protein also makes contact with the μ1 protein that forms a lattice on the surface of the ISVPs (26). Because the λ2 protein is an intermediary between the σ1 and μ1 proteins (26), a change in the σ1-λ2 interaction could transduce a signal to μ1 and alter its properties. Since both the attachment and ISVP* phenotype of T3D^F^/T3D^C^S1 could be explained by σ1-λ2 interaction, we decided to determine whether a mismatch between T3D^C^ σ1 and T3D^F^ λ2 accounts for the phenotypes observed. The λ2 proteins of T3D^F^ and T3D^C^ differ at residue 504 resulting in a Glycine-to-Glutamate change and at residue 509 resulting in a Glycine-to-Arginine change (69). Thus, to test our idea about the contribution of σ1-λ2 mismatch, we generated two additional viruses, T3D^F^/T3D^C^L2 (a monoreassortant that contains a L2 gene segment from strain T3D^C^ in an otherwise T3D^F^ background) and T3D^F^/T3D^C^S1L2 (a double reassortant that contains both the S1 and L2 gene segments from strain T3D^C^ in an otherwise T3D^F^ background). Among these, σ1 and λ2 remain mismatched in T3D^F^/T3D^C^L2 whereas they are matched in T3D^F^/T3D^C^S1L2. In comparison to T3D^F^, plaques formed by T3D^F^/T3D^C^L2 and T3D^F^/T3D^C^S1L2 were predominantly smaller. Notably, the plaque size of both viruses and most notably T3D^F^/T3D^C^S1L2 were larger than those formed by T3D^F^/T3D^C^S1 (Fig 6A, compare to Fig 1A).

**Figure 6.**
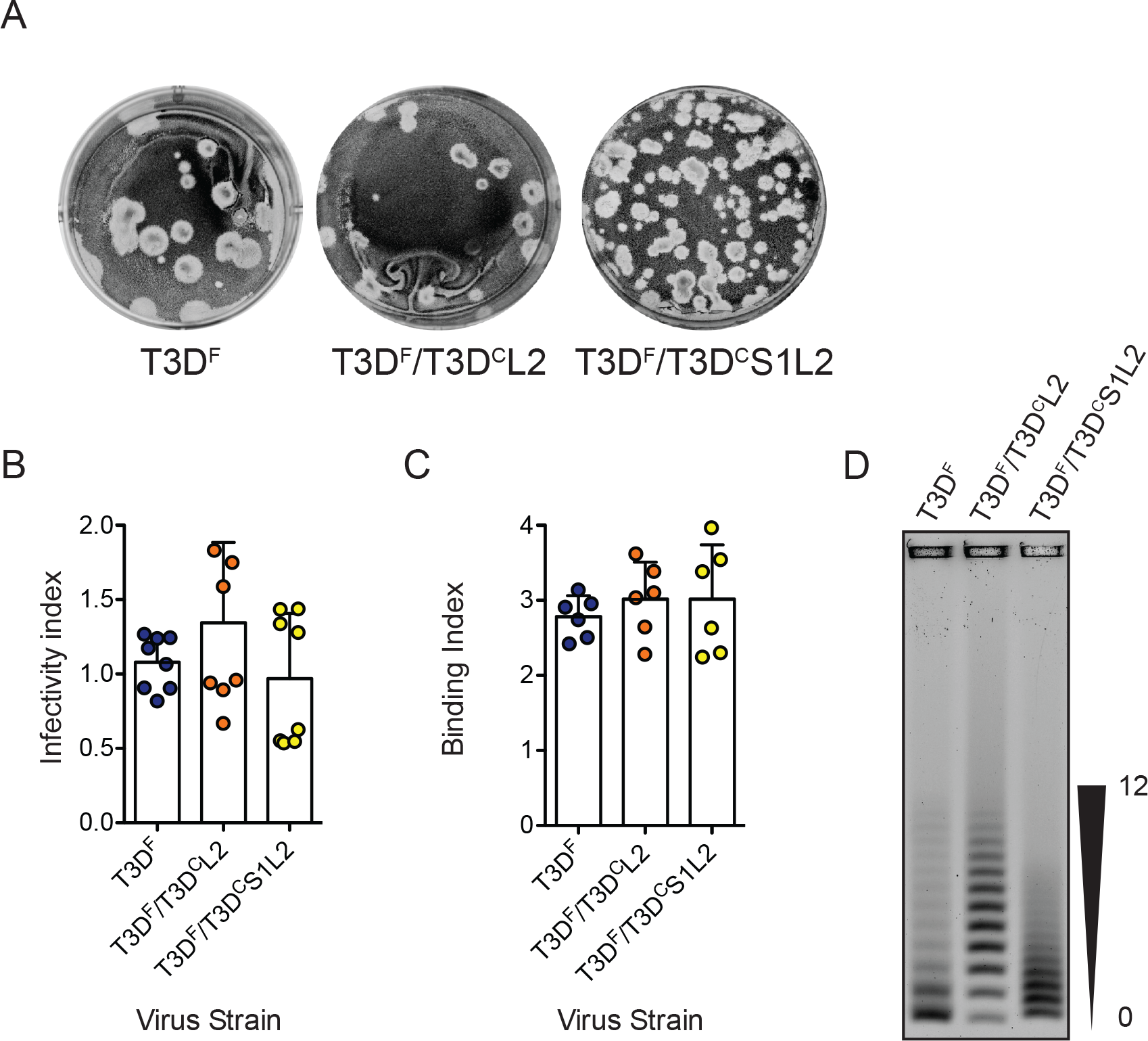
Matched L2 gene from T3D^C^ restores σ1 encapsidation defect in T3D^F^/T3D^C^S1. (A) Particles of either T3D^F^, T3D^F^/T3D^C^L2 or T3D^F^/T3D^C^S1L2 diluted in PBS were subjected to plaque assay on spinner adapted L929 cells. This plaque assay was performed in parallel with one shown in Fig 1 (B) ATCC L929 cells were adsorbed with 3000 particles/cell of either T3D^F^, T3D^F^/T3D^C^L2 or T3D^F^/T3D^C^ S1L2. After incubation at 37°C for 18 h, the cells were subjected to indirect immunofluorescence assay using a LI-COR Odyssey scanner. Relative infectivity was determined by calculating intensity ratios at 800 nm (green fluorescence) representing viral antigen and 700 nm (red fluorescence) representing the cell monolayer. Infectivity index for each independent infection and the sample mean are shown. Error bars indicate SD. NS, P > 0.05 as determined by 1-way ANOVA with Bonferroni’s multiple comparison test in comparison to T3D^F^ (C) Confluent monolayers of L929 cells grown in 96-well plates were adsorbed with 5 × 10^4^ particles/cell of either T3D^F^, T3D^F^/T3D^C^L2 or T3D^F^/T3D^C^S1L2 at 4°C for 1 h. Cell attachment was determined by indirect immunofluorescence of cell associated particles using a LI-COR Odyssey scanner. Binding index was determined by calculating intensity ratios at 800 nm (green fluorescence) representing viral antigen and 700 nm (red fluorescence) representing the cell monolayer. Binding index for each independent infection and the sample mean are shown. Error bars indicate SD. NS, P > 0.05 as determined by 1-way ANOVA with Bonferroni’s multiple comparison test in comparison to T3D^F^ (D) Virions (1 × 10^11^ particles) were resolved on an agarose gel, stained with colloidal blue staining kit and scanned using a LI-COR Odyssey scanner. The position of virions with lowest and highest number of σ1 trimers is shown.

To further evaluate the phenotype of T3D^F^/T3D^C^L2 and T3D^F^/T3D^C^S1L2, we infected L929 cells with 3000 particles/cell of each virus and assessed their infectivity using indirect immunofluorescence. We found that T3D^F^/T3D^C^L2 and T3D^F^/T3D^C^S1L2 established infection as efficiently as T3D^F^ (Fig 6B). These data indicate that changing the L2 sequence through reassortment of a T3D^C^ L2 gene alone does not influence the capacity of virus to establish infection. Additionally, based on the comparison between the infectivity of T3D^F^/T3D^C^S1 (Fig 1) and T3D^F^/T3D^C^S1L2, our data indicate that introduction of a matched T3D^C^ L2 gene segment into T3D^F^/T3D^C^S1 restores the infectivity of the virus to wild-type level.

Lower infectivity of T3D^F^/T3D^C^S1 is a consequence of reduced capacity of this virus strain to attach host cells. To examine the cell attachment efficiency of T3D^F^/T3D^C^L2 and T3D^F^/T3D^C^S1L2, we used a plate-based quantitative attachment assay. We found that T3D^F^/T3D^C^L2 and T3D^F^/T3D^C^S1L2 displayed attachment comparable to T3D^F^ (Fig 6C). These data indicate that the attachment phenotype produced by the introduction of T3D^C^ σ1 into T3D^F^, is overcome by simultaneous introduction of a matched T3D^C^-derived λ2 protein.

Based on the attachment properties of T3D^F^/T3D^C^L2 and T3D^F^/T3D^C^S1L2, it was expected that virions of this strain would exhibit T3D^F^-like σ1 encapsidation. To directly test this possibility, purified virus particles were resolved on agarose gels. We observed that in comparison to T3D^F^, the pattern of distribution of viruses based on σ1 trimer encapsidation was different (Fig 6D). The T3D^F^/T3D^C^L2 preparation contained a higher proportion of virions with more σ1. Because of the efficient encapsidation of σ1 on the particle, T3D^F^/T3D^C^L2 is able to efficiently attach and infect host cells. The preparation of T3D^F^/T3D^C^S1L2 contained virions with varied levels of σ1 trimers. Though the T3D^F^/T3D^C^S1L2 preparation does not contain a large proportion of virions with higher number of σ1 trimers observed for T3D^F^, it contains sufficient number of particles with the needed level of σ1 trimers to allow T3D^F^-like infection. Importantly, despite the shared T3D^C^ S1 allele, the presence of a large proportion of virions with no σ1 trimers noted for T3D^F^/T3D^C^S1 is not observed in the preparation of T3D^F^/T3D^C^S1L2 (compare to Figure 3). These data indicate that both the nature of σ1 and λ2 influence the encapsidation of σ1 trimers on the particle. Furthermore, these data indicate that the encapsidation defect produced by reassortment of the T3D^C^ S1 into T3D^F^ can be overcome by the presence of a T3D^C^-derived matched λ2 protein.

### Properties of the λ2 protein influence ISVP-to-ISVP* conversion

In addition to the effect of T3D^C^ S1 on attachment and infection, we also observed that in comparison to T3D^F^, T3D^F^/T3D^C^S1 forms ISVP*s much more readily, under less stringent conditions (Figure 5). Based on the evidence that inclusion of T3D^C^ λ2 restored the attachment phenotype of T3D^F^/T3D^C^S1, we determined the capacity of T3D^F^/T3D^C^L2 and T3D^F^/T3D^C^S1L2 to undergo ISVP-to-ISVP* conversion. Chymotrypsin-generated ISVPs of each virus were incubated at increasing temperature and the conditions under which the δ fragment of μ1 attains a trypsin-sensitive conformation was determined. We found that in comparison to T3D^F^ ISVPs, which display a trypsin-sensitive conformer of δ at 42°C (Fig 7A), the T3D^F^/T3D^C^L2 5 fragment attains a trypsin-sensitive conformation at a lower temperature of 32°C (Fig 7B). These data indicate that properties of the λ2 protein influence the propensity with which ISVP*s are generated. The temperature at which T3D^F^/T3D^C^S1L2 converted to ISVP* was higher (34°C) than that of T3D^F^/T3D^C^L2 (Fig 7C). Importantly, though T3D^F^/T3D^C^S1L2 converted to ISVP* in a more facile manner than the parental T3D^F^ strain, its propensity to form ISVP*s was lower than that of mismatched viruses, T3D^F^/T3D^C^S1 or T3D^F^/T3D^C^L2, which convert to ISVP* at ~ 28°C and 32°C respectively (compare to Fig 5). These data indicate that placement of the T3D^C^ L2 in T3D^F^/T3D^C^S1 raises the temperature at which ISVPs undergo ISVP-to-ISVP* conversion. To assess ISVP* formation by monitoring loss of infectivity of ISVPs, in vitro generated ISVPs of T3D^F^, T3D^F^/T3D^C^L2 and T3DF/T3D^C^S1L2 were heated at increasing temperatures and the infectivity of the samples on L929 cells was monitored using the quantitative indirect immunofluorescence assay. We observed that while T3D^F^ lost infectivity at 42°C, T3D^F^/T3D^C^L2 and T3D^F^/T3D^C^S1L2 lost infectivity at 30°C and 32°C respectively (Fig 7D, 7E and 7F). Importantly, the temperature at which T3D^F^/T3D^C^S1L2 loses infectivity is higher than that for T3DF/T3D^C^S1 and T3D^F^/T3D^C^L2 (28°C and 30°C respectively) (Fig 5). Thus, our results highlight a previously unknown effect of λ2 on the efficiency of ISVP-to-ISVP* conversion. Further, our data also indicate that a match between the σ1 and λ2 proteins controls conformational changes in virus capsid that are required for bypassing cellular membranes during cell entry.

**Figure 7.**
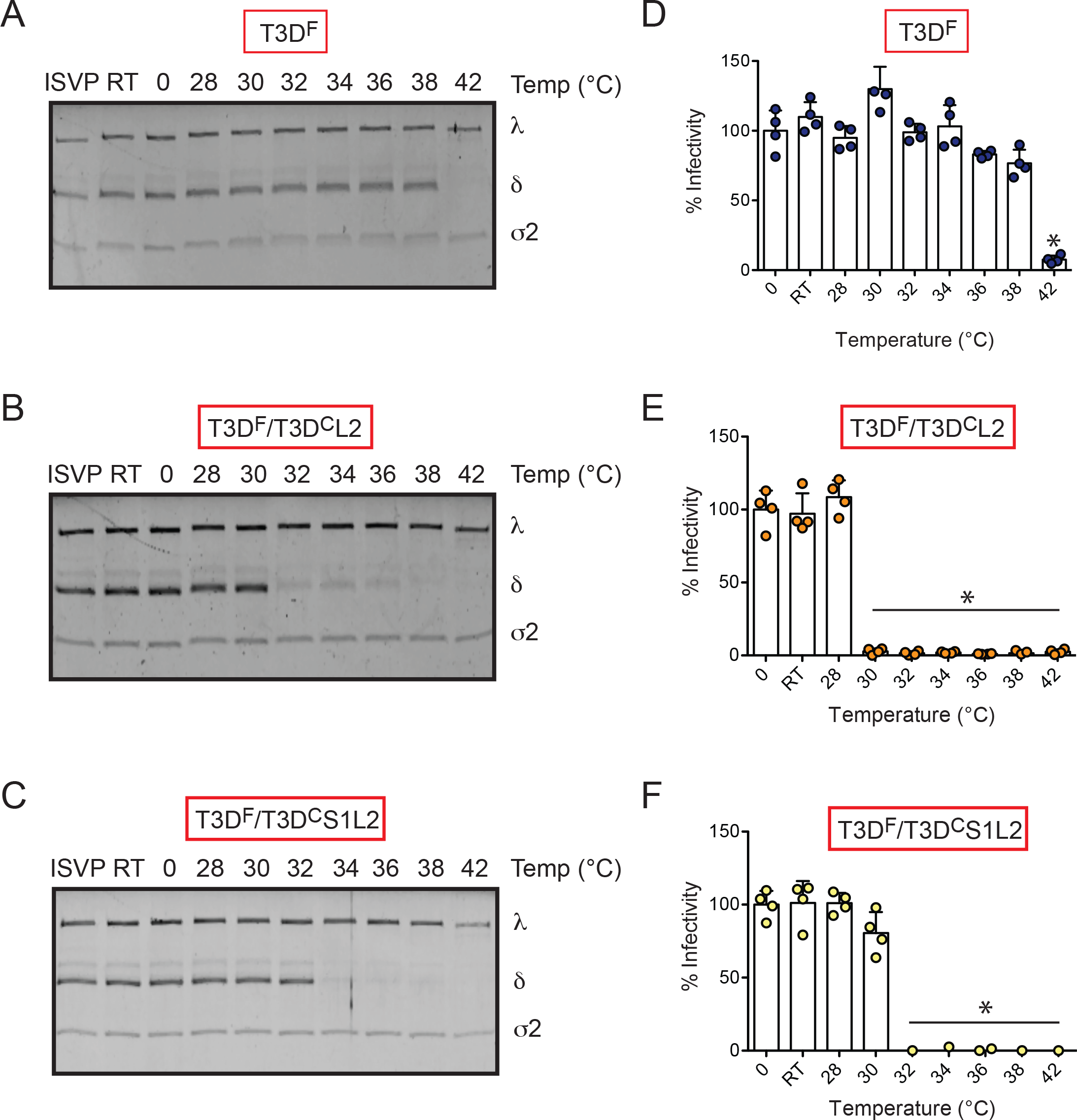
The *λ2* protein influences ISVP-to-ISVP* conversion. (A, B, C) ISVPs (2 × 10^12^ particles/ml) of T3D^F^, T3D^F^/T3D^C^L2 or T3D^F^/T3D^C^S1L2 were divided into aliquots of equivalent volume and heated at temperatures ranging from 0°C to 42°C for 20 min. The reactions were chilled on ice and then digested with 0.10 mg/ml trypsin for 30 min on ice. Following addition of loading dye, the samples were subjected to SDS-PAGE analysis. The gels shown are representative of at least 3 independent experiments (D, E, F) ISVPs of (2 × 10^12^ particles/ml) of T3D^F^, T3D^F^/T3D^C^L2 or T3D^F^/T3D^C^S1L2 were heated at 0°C to 42°C for 20 min and then used to initiate infection of L929 cells at 300 particles/cell. After incubation at 37°C for 18 h, the cells were subjected to indirect immunofluorescence assay using a LI-COR Odyssey scanner. Relative infectivity was determined by calculating intensity ratios at 800 nm (green fluorescence) representing viral antigen and 700 nm (red fluorescence) representing the cell monolayer. Infectivity index of ISVPs maintained at 0°C was set to 100%. Percent infectivity for each independent infection and the sample mean are shown. Samples with a calculated infectivity below 0% are not shown. Error bars indicate SD. *, P < 0.05 as determined by 1-way ANOVA with Bonferroni’s multiple comparison test in comparison to T3D^F^.

## DISCUSSION

To test the idea that new protein-protein interactions forced by placement of alleles from two parent viruses in a reassortant virus alter the infectious properties of virus capsids, we compared the T3D^F^ parent with T3D^F^/T3D^C^S1, a monoreassortant containing the S1 gene from laboratory isolate T3D^C^. In comparison to T3D^F^, T3D^F^/T3D^C^S1 encapsidates the S1-encoded σ1 protein less efficiently, resulting in a population of particles that have no σ1. Given the function of σ1 in attachment, this defect results in a lower capacity of T3D^F^/T3D^C^S1 to attach and infect host cells. Co-introduction of the T3D^C^-derived L2 gene, which encodes the λ2 protein that anchors fibers of the σ1 protein into the virion, allowed better encapsidation of σ1 and restored the attachment and infectivity defect observed for T3D^F^/T3D^C^S1. These data indicate that reassortment could influence how adjacent capsid proteins fit with each other and consequently influence the function of the capsid.

The σ1 protein folds into a homotrimeric structure with a distinct head, body, and tail domain (70, 71). The N-terminal end of the protein is predicted to form an α-helical coiled coil (70, 71). This region transitions into a body domain that is formed from β-spiral repeats (72). The C-terminal portion of the protein forms a globular head structure with an eight-stranded β-barrel (72). σ1 is anchored to the particles via turrets formed by the λ2 protein (26, 68). While structural evidence highlighting contacts between σ1 and λ2 is lacking, mutational analyses indicate that a σ1 region comprised of amino acids 334, which includes a hydrophobic stretch, is necessary for anchoring into λ2 (73, 74). The T3D^F^-T3D^C^ polymorphism at residue 22 in σ1 produces a Valine-to-Alanine change, which could alter the hydrophobicity of the σ1 N-terminus and influence its encapsidation on the particle. The second T3D^F^-T3D^C^ polymorphism at residue 408 lies within the head domain of σ1. This region is not thought to engage any portion of the virus and thus not expected to directly influence encapsidation of σ1 on the virus particle. Our results indicate that the attachment defect produced by the presence of the T3D^C^ σ1 protein is overcome by introduction of the T3D^C^ λ2 protein. Though σ1 encapsidation on T3D^F^/T3D^C^L2 and T3D^F^/T3D^C^S1L2 is not identical to that observed for T3D^F^ (Figure 6), this difference does not appear to influence the capacity of the virus to attach and infect cells. This result is consistent with previous observations indicating that a threshold number of σ1 on particles is sufficient for the virus to establish infection and that particles with less σ1 than the parental strain retain their infectivity (58, 59).

The λ2 protein forms a pentameric turret at the 5-fold vertices of the virus (21). The hollow cavity formed by λ2 is lined by enzymatic domains that are required for capping viral mRNA (21). Additionally, in virions and ISVPs, the λ2 pentamer holds the σ1 fiber via the flaps formed by the last 250 amino acid residues of λ2 (21, 26, 41). Thus, the T3D^C^ σ1 encapsidation defect may be a consequence of a mismatch between T3D^C^ σ1 and T3D^F^ λ2. The amino acid sequence of T3D^F^ and T3D^C^ λ2 proteins differs at two residues, 504 and 509, that constitute the methylase-1 domain (21). No changes are present in the flap region of λ2 that is proposed to interact with σ1. Thus, why an allelic match between σ1 and λ2 is required for optimal attachment is not immediately obvious. The arginine at position 509 is in a location to be able to make interactions with the underside of the flap domain comprised of residues 1263-1267 (Fig 8). Such an interaction may alter the conformation of the λ2 flaps and alter its capacity to assemble or retain σ1 efficiently. Our observations with T3D^F^/T3D^C^L2 indicate that even if changes at residues 504 and 509 do influence the local structure of λ2, its effects are only manifested in the presence of T3D^C^ σ1. Whether this is due to the greater hydrophobicity of the T3D^F^ σ1 protein or some other effects is not known. It should be noted that even though the parental virus T3D^C^ contains a matched σ1-λ2 pair, similar to T3D^F^/T3D^C^S1L2, it encapsidates more σ1 than T3D^F^ (36). Thus, σ1 encapsidation may also be influenced by the properties of other capsid proteins. Indeed, we have previously observed that a μ1 variant contains different levels of particle-associated σ1 (75). Another study has also suggested that compatibility between μ1 and σ1 confers optimal σ1 encapsidation and function on the particle (69). The difference in σ1 encapsidation between T3D^F^ and T3D^C^ may be governed by the function of μ1 or yet another protein.

**Figure 8.**
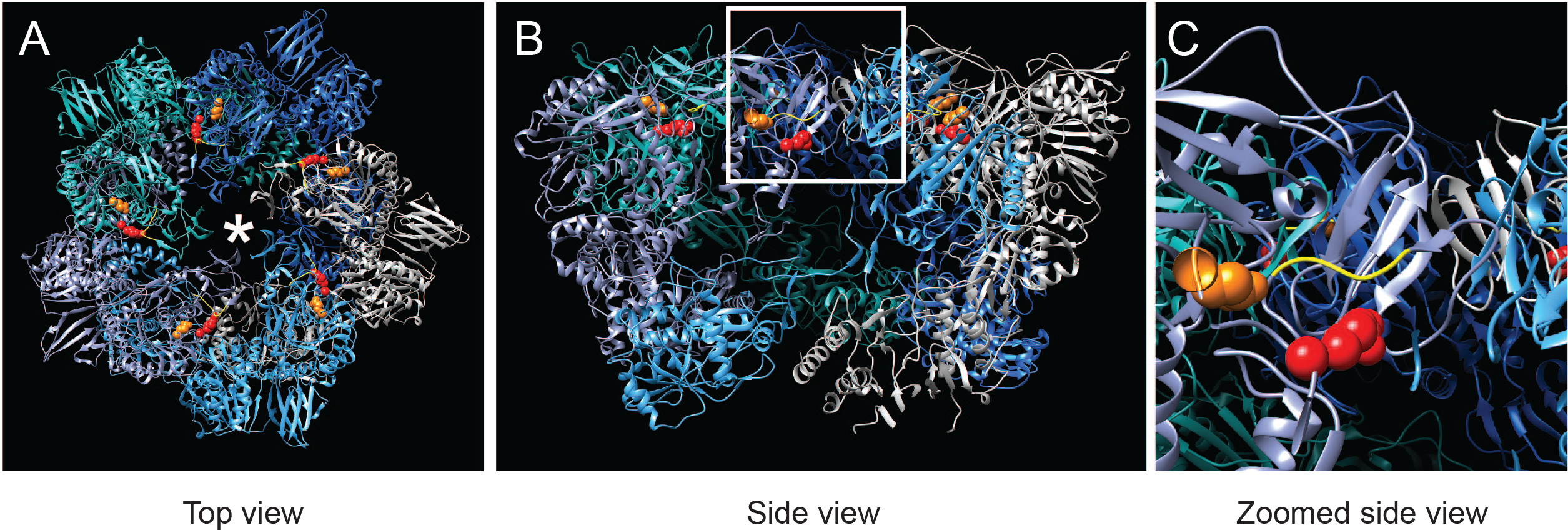
Position of λ2 polymorphisms on λ2 pentamer. (A) Top view of the λ2 pentamer rendered using UCSF Chimera from PDB 2CSE is shown with each λ2 monomer in a different color (91). The positions of residue 504 (orange), and 509 (red) and the σ1 trimer are shown using an asterix (*). The yellow ribbon indicates the position of the underside of the λ2 flap domain. (B, C) A side of the same molecule (B) and zoomed in inset are shown (C).

Capsids of T3D^F^ and T3D^F^/T3D^C^S1 differ in at least one unexpected way. In comparison to T3D^F^, T3D^F^/T3D^C^S1 undergoes ISVP-to-ISVP* conformational changes that are required for cell entry more readily. As described above, the only known interaction between σ1 and the particle is via the λ2 pentamer (26, 68). As a consequence of ISVP-to-ISVP* transition, λ2 adopts an open conformation similar to the one found in core particles, consequently allowing the release of the σ1 protein and the derepression of viral transcriptase activity (27). The δ fragment derived from the μ1 protein also undergoes a structural transition during ISVP* formation, generating a protease-sensitive conformer (27, 76). The outer side of the λ2 turret contacts δ in ISVP particles (26). Thus, changes to structure of λ2 likely influence its interaction with δ. It is possible that changes to the structure of each of the proteins is hierarchical with one structural change being a prerequisite for another. Alternatively, the different components of the capsid may function like an interlocking gear set where changes to one protein (for example λ2) cannot occur without a concurrent change to another part of the capsid (for example δ). We hypothesize the presence of σ1 serves as a lock such that a lower level of this protein in the λ2 pentamer or one that is loosely held (such as in T3D^F^/T3D^C^S1) would favor conformational changes in the remaining capsid. Our work using T3D^F^/T3D^C^L2 and T3D^F^/T3D^C^S1L2 indicates that a virus that bears matching T3D^C^ σ1 and λ2 proteins displays a more regulated conversion to ISVP* than either mismatched virus. Because the level of σ1 on T3D^F^/T3D^C^L2 and T3D^F^/T3D^C^S1L2 are higher than those on T3D^F^/T3D^C^S1, the level of σ1 does not appear to correlate with the capacity of the virus to form ISVP*. Based on our characterization of T3D^F^/T3D^C^L2, our studies also revealed for the first time that the nature of the λ2 influences the efficiency of ISVP-to-ISVP* conversion. As stated above, the amino acid residues that are different between T3D^F^ and T3D^C^ λ2 are proximal to the inner cavity of the λ2 pentamer proteins and are not in position to influence interaction with δ on the outer side of λ2. Thus, one possibility is that there is a significant structural difference in the T3D^F^ and T3D^C^ λ2 pentamer. We however favor a different possibility to explain our results. This idea is an extension of our hypothesis about the locking structures in the virus. Reovirus ISVPs are metastable structures that are poised to undergo structural changes (27, 67). The metastability of ISVPs is maintained by interactions between capsid proteins (42, 77, 78). When such interactions are altered by placing alleles that are not coevolved together, it renders particles more metastable and thus more likely to readily convert to ISVP*. We think this is an underappreciated effect of reassortment.

Following intranasal inoculation, in comparison to T3D^F^, T3D^C^ more efficiently replicates in the lungs and disseminates to secondary sites (36). Based on the evidence that T3D^F^/T3D^C^S1 replicates more efficiently at secondary sites of infection than T3D^F^, this difference was attributed to the S1 gene segment and a role for both σ1 and σ1s was described (36). T3D^C^ incorporates significantly more σ1 than T3D^F^ (36). However, based on our evidence that T3D^F^/T3D^C^S1 incorporates less σ1 than T3D^F^, we think it is unlikely that the level of σ1 controls the difference in the in vivo properties of T3D^F^ and T3D^C^ or T3D^F^/T3D^C^S1. In comparison to T3D^F^, T3D^F^/T3D^C^S1 is less sensitive to inactivation by proteases (36). Because how this compares to T3D^C^ is not known, the contribution of protease sensitivity to more superior replication of T3D^F^/T3D^C^S1 in vivo is undefined. Why the lower level of σ1 incorporation does not compromise replication of T3D^F^/T3D^C^S1 in vivo is puzzling. We speculate that differences in the level of host cell receptors in different tissues accessed by the virus following intranasal inoculation or the use of alternate receptors such as Ngr1, which may engage parts of the virus other than σ1 may allow the virus to replicate efficiently in vivo (79). Our previous work indicates that mutants that are impaired in the capacity to convert to ISVP*s are compromised for replication at least in the central nervous system (80). Whether replication in other tissues is influenced by ISVP-to-ISVP* conversion efficiency remains to be determined. Determinants of viral replication in different mouse tissues are different and it is possible that the impact of this stage of virus replication is only evident in some tissues (81–84).

## MATERIALS AND METHODS

### Cells

Spinner-adapted Murine L929 cells were maintained in Joklik’s MEM (Lonza) supplemented to contain 5% fetal bovine serum (FBS) (Sigma-Aldrich), 2 mM L-glutamine (Invitrogen), 100 U/ml penicillin (Invitrogen), 100 μg/ml streptomycin (Invitrogen), and 25 ng/ml amphotericin B (Sigma-Aldrich). Spinner-adapted L929 cells were used for cultivating, purifying and titering viruses. L929 cells obtained from ATCC were maintained in Eagle’s MEM (Lonza) supplemented to contain 5% fetal bovine serum (FBS) (Sigma-Aldrich), and 2 mM L-glutamine (Invitrogen). Experiments to measure attachment and infectivity were done using L929 cells from ATCC.

### Generation of Recombinant Viruses

Changes to the sequence of pT7-T3D S1 and pT7-T3D L2 were engineered using QuikChange lightning site-directed mutagenesis kit (Agilent). Plasmid sequences were confirmed by sequencing. Recombinant strains T3D^F^/T3D^C^S1 which contains T3D^F^ S1 gene with mutations (V22A and T408A), T3D^F^/T3D^C^L2 which contains a mutated T3D^F^ L2 gene (G504E and G509R) and T3D^F^/T3D^C^S1L2 which contains a T3D^F^ S1 gene with mutations (V22A and T408A) with mutated T3D^F^ L2 gene (G504E and G509R) in an otherwise T3D^F^ background were generated by using a plasmid-based reverse-genetics strategy (85, 86). To confirm sequences of mutant viruses, viral RNA was extracted from infected cells or virus particles and subjected to reverse transcription PCR using S1-specific or L2-specific primers. PCR products were resolved on Tris-acetate-EDTA agarose gels, purified and confirmed by sequence analysis. Primer sequences will be shared upon request.

### Purification of viruses

Purified reovirus virions were generated using second - or third-passage L-cell lysates stocks of reovirus. Viral particles were Vertrel-XF (Dupont) extracted from infected cell lysates, layered onto 1.2- to 1.4-g/cm^3^ CsCl gradients, and centrifuged at 187,183 × *g* for 4 h. Bands corresponding to virions (1.36 g/cm^3^) were collected and dialyzed in virion-storage buffer (150 mM NaCl, 15 mM MgCl2, 10 mM Tris-HCl [pH 7.4]) (63). The concentration of reovirus virions in purified preparations was determined from an equivalence of one OD unit at 260 nm equals 2.1 × 10^12^ virions/ml (87). Virus titer was determined by plaque assay on spinner-adapted L929 cells. At least three preparations each of T3D^F^, T3D^F^/T3D^C^S1, T3D^F^/T3D^C^L2 and T3D^F^/T3D^C^S1L2 were used for this study.

### Assessment of Plaque Morphology

Plaque assays were conducted in spinner-adapted L929 cells plated in 6-well plates (Greiner Bio-One). Cells were adsorbed with dilutions of virus in PBS. Cells were overlaid with a molten mixture comprised of 1X Medium 199 and 1% Bacto agar supplemented with 10 μg/ml chymotrypsin. 7 days following infection, the monolayers were fixed by addition of 4% Formaldehyde solution in PBS and incubated overnight. The agar overlay was peeled off and the monolayers were stained with 1% Crystal violet stain in 5% ethanol for 5 hours at room temperature. The monolayers were washed with water. Plaque sizes were noted by visual comparison.

### Generation of ISVPs in vitro

ISVPs were generated in vitro by incubation of 2 × 10^12^ virions with 200 μg/ml of TLCK-treated CHT at 32°C in virion storage buffer (150 mM NaCl, 15 mM MgCl_2_, 10 mM Tris-HCl [pH 7.4] for 20 or 60 min. Proteolysis was terminated by addition of 2 mM phenylmethylsulphonyl fluoride (PMSF) and incubation of reactions on ice. Generation of ISVPs was confirmed by SDS-polyacrylamide gel electrophoresis (PAGE) and Coomassie Brilliant Blue staining.

### Assessment of infectivity by indirect immunofluorescence

Monolayers of L929 cells (4 × 10^4^) in clear bottom tissue culture treated 96-well plates (Corning) were washed with PBS and infected with virions or ISVPs of the indicated reovirus strain at 4°C for 1 h. In experiments to examine the role of glycans on infection, the cells were pretreated with serum free Eagle’s MEM containing 10 mU/ml Neuraminidase (Roche) at 37°C for 1 h prior to virus attachment. In experiments to examine the effect of 9BG5 mAb on virus infectivity, 1 × 10^11^ virus particles/ml were incubated at 4°C with 500 ng/ml of hybridoma supernatant containing 9BG5 mAb overnight (54, 88). This mixture was used to initiate infection as described above. Monolayers were fixed with methanol at −20°C for a minimum of 30 min and washed with PBS containing 0.5%Tween-20 (DPBS-T). Cells were then incubated with polyclonal rabbit anti-reovirus serum at a 1:1000 dilution in PBS with 1% BSA (DPBS-BSA) at 37°C for 60 min (89). Monolayers were washed twice with DPBS-T and incubated with DPBS-BSA for 37°C for 60 min followed by two washes with DPBS-T. Cells were stained with 1:1000 dilution of LI-COR CW 800 anti-rabbit immunoglobin G, 1:1000 dilution of Sapphire 700 (LI-COR) and DRAQ5 (Cell signaling Technology) at a concentration of 1:10000 for 37°C for 60 min. Monolayers were washed thrice with DPBS-T and fluorescence intensity was measured using the Odyssey Imaging System and the Image Studio Lite software (LI-COR). For each well, the ratio of fluorescence at 800 nm (for infected cells) and 700 nm (for total cells) was quantified. Relative infectivity in arbitrary units was quantified using the following formula. Relative Infectivity = (Green/Red)_infected_ - (Green/Red)_uninfected_.

### Assessment of infectivity of input and progeny virus

In experiments to determine infectivity of virus released from infected cells, L929 cells were adsorbed with virions at room temperature for 1 h. After incubation at 37°C for 24 h, 25 μ1 of the supernatant from infected wells was used to initiate infection of fresh L929 cells plated in 96-well plates. Plate 1 was fixed with cold methanol. The infection in Plate 2 was allowed to proceed at 37°C for 24 h. Infectivity was measured using indirect fluorescence assay described above.

### Assessment of reovirus attachment

L929 cells (4 × 10^4^ cell per well) grown in 96-well plates were chilled at 4°C for 15 min and then adsorbed with particles of the indicated virus strains at 4°C for 1 h. Cells were washed with chilled PBS and blocked with PBS-BSA at 4°C for 15 min. Cells were then incubated with polyclonal rabbit anti-reovirus serum at a 1:2500 dilution in PBS-BSA at 4°C for 30 min. The cells were washed twice with PBS-BSA followed by incubation with 1:1000 dilution of Alexa Fluor 750 labeled goat anti-rabbit antibody at 4°C for 30 min. After two washes with PBS-BSA, cells were stained with 1:1000 dilution of DNA stain, DRAQ5 (Cell Signaling technology), at 4°C for 5 min. Cells were washed and then fixed with 4% formaldehyde at room temperature for 20 min. Fluorescence intensity was measured using the Odyssey Imaging System and the Image Studio Lite software (LI-COR). For each well, the ratio of fluorescence at 800 nm (for attached reovirus) and 700 nm (for total cells) was quantified. Binding index in arbitrary units was quantified using the following formula. Binding index = (Green/Red)_infected_ - (Green/Red)_uninfected_.

### Analysis of protein levels by immunoblotting

The samples: purified virions or whole cell lysates of infected cells prepared using RIPA lysis buffer (50 mM NaCl, 1 mM EDTA at pH 8, 50 mM TRIS at pH 7.5, 1% Triton X-100, 1% sodium deoxycholate, 0.1% SDS) supplemented with protease inhibitor cocktail (Roche) and 500 μM PMSF were resolved on 10% SDS-PAGE gels and transferred to nitrocellulose membranes. For immunoblots using polyclonal rabbit anti-reovirus serum, the membranes were blocked with 5% milk in TBS at room temperature for 1 h. For immunoblots using 4A3 mouse anti-μ1 mAb and rabbit anti T3D σ1 head antibody, membranes were blocked with T20 Starting Block (Thermofisher Scientific)(54, 90). Following blocking, rabbit anti-reovirus serum (1:1000), 4A3 mouse anti-μ1 mAb (1:500), and rabbit anti T3D σ1 head antibody (1:750) were incubated with the membrane in appropriate blocking buffer at room temperature for 1 h. The membranes were washed with TBS supplemented with 0.1% Tween 20 (TBS-T) twice for 15 min and then incubated with Alexa Fluor conjugated anti-rabbit IgG or anti-mouse IgG in blocking buffer. Following three washes, membranes were scanned using an Odyssey Infrared Imager (LI-COR) and intensities of μ1C and σ1 bands intensity were quantified using the Image studio lite software (LI-COR).

### Hemagglutination (HA) assay

Purified T3D^F^ or T3D^F^/T3D^C^S1 virions (1 × 10^11^ particles) were serially diluted in 50 μl of PBS in 96-well V bottom microtiter plates (Corning-Costar). Bovine erythrocytes (Colorado Serum Company) were washed twice with chilled PBS and were resuspended at a concentration of 1% (vol/vol) in PBS. Washed erythrocytes (50 μl) were added to wells containing virus and incubated at 4°C overnight. HA titer was expressed as 2 × 10^11^ particles divided by the number of particles per HA unit. One HA unit is equal to the number of particles of virus sufficient to produce HA (50).

### Agarose gel separation of reovirus particles by σ1 content

1 × 10^11^ virus particles were resuspended in dialysis buffer, mixed with 2X Gel loading dye (NEB) and resolved on 1% ultra-pure agarose gel (Invitrogen) in 1X TAE pH 7.2 at constant 25 V for 18 h. The gel was stained with NOVEX colloidal blue staining kit (Invitrogen) for 6 h and destained overnight in water. Gel was scanned using Odyssey Infrared Imager (LI-COR). Distribution of virions with different σ1 levels was interpreted by visual comparison.

### Analysis of ISVP-to-ISVP* conversion

ISVPs (2 × 10^12^ particles/ml) of the indicated viral strains were divided into aliquots of equivalent volume and heated at temperatures ranging from 0°C to 42°C for 20 min. The reactions were cooled on ice and then digested with 0.10 mg/ml trypsin (Sigma-Aldrich) for 30 min on ice. Following addition of the SDS PAGE loading dye, the samples were subjected to SDS-PAGE analysis. For analysis by quantitative infectivity assay, the heated samples were used to initiate infection of L929 cells.

### Statistical analysis

Statistical significance between two experimental groups was determined using the Student’s t-test function of the GraphPad Prism software. When an experimental group was compared to more than one other experimental group, 1-way ANOVA with Bonferroni’s multiple comparison test function of the GraphPad Prism software was used.

## ACKNOWLEDGEMENTS

We thank Karl Boehme, Tuli Mukhopadhyay, John Patton, and Anthony Snyder along with members of our laboratory for helpful suggestions and review of the manuscript.

